# Mutations in the juxtamembrane segment of the cholesterol-binding site of APP alter its processing and promotes production of shorter, less toxic Aβ peptides

**DOI:** 10.1101/2020.11.16.384891

**Authors:** Linda Hanbouch, Béatrice Schaack, Amal Kasri, Gaëlle Fontaine, Eleni Gkanatsiou, Gunnar Brinkmalm, Erik Portelius, Kaj Blennow, Gilles Mourier, Nicolas Gilles, Mark J Millan, Catherine Marquer, Henrik Zetterberg, Lydie Boussicault, Marie-Claude Potier

## Abstract

**Background:** The brains of patients with Alzheimer’s disease (AD) reveal increased cellular membrane levels of cholesterol. Correspondingly, we previously showed that elevating levels of membrane cholesterol in neuronal cultures recapitulates early AD phenotypes including excessive cleavage of amyloid β (Aβ) peptides from the amyloid precursor protein (APP). Here we aimed to evaluate how the presence of a cholesterol-binding site (CBS) in the transmembrane and juxtamembrane regions of APP regulates its processing.

**Methods:** We generated seven single and two double APP mutants at amino acid positions 22, 26, 28, 29, 33, 39 of the Aβ sequence changing the charge and/or hydrophobicity of the targeted amino acids. HEK293T cells were transfected with APP constructs and secreted Aβ peptides were measured using ELISA and mass spectrometry (MS). APP processing in normal and high cholesterol condition, and endocytosis were assessed in stably expressing APP^wt^ and APP^K28A^ HEK293T clones. Finally, we measured the binding of synthetic peptides derived from the Aβ sequence to cholesterol-rich exosomes purified from control HEK293T cells.

**Results:** Most mutations triggered a reduction in the production of Aβ40 and Aβ42 peptides, whereas only juxtamembrane mutants resulted in the generation of shorter Aβ peptides. We confirmed by mass spectrometry this specific change in the profile of secreted Aβ peptides for the most characteristic APP^K28A^ mutant. A transient increase of plasma membrane cholesterol enhanced the production of Aβ40 by APP^WT^, an effect absent with APP^K28A^. The enzymatic activity of α-, β- and γ-secretases remained unchanged in cells expressing APP^K28A^. Similarly, APP^K28A^ subcellular localization in early endosomes did not differ to APP^WT^. Finally, WT but not CBS mutant Aβ derived peptides bound to cholesterol-rich exosomes.

**Conclusions:** Taken together, these data reveal a major role of the juxtamembrane region of APP in binding to cholesterol and accordingly in the regulation of APP processing. Binding of cholesterol to K28 could staple APP to the juxtamembrane region thereby permitting access to γ-secretase cleavage at positions 40-42. The APPK28 mutant would lie deeper in the membrane, facilitating the production of shorter Aβ peptides and unveiling this specific region as a novel target for reducing the production of toxic Aβ species.

## Background

Alzheimer’s disease (AD) is the most common form of dementia in the elderly population and is characterized by two prominent pathologies, extracellular amyloid-β (Aβ) containing plaques and intraneuronal fibrillary tangles comprised of aberrantly hyperphosphorylated tau protein (1). Aβ peptides of various lengths are produced by sequential proteolysis of the transmembrane Amyloid Precursor Protein (APP) by the β-secretase BACE1 and the γ-secretase which operate in the membrane bilayer (2). Amyloidogenic APP processing occurs in the endolysosomal compartment following clathrin-dependent APP internalization (3).

There is considerable interest in endogenous factors controlling the processing of APP and their potential therapeutic modulation. One line of research has focused on cholesterol, which is produced in the brain (independently of the periphery) by astrocytes, then shuttled to neurons bound to apolipoprotein E (APOE) protein. APOE encoded by the polymorphic gene *APOE* which possesses three alleles ε2, ε3 and ε4 (4), with the strongest genetic risk factor for sporadic AD being the ε4 allele of *APOE* (5). Levels of cholesterol are elevated in the brain of people diagnosed with AD and it is known to accumulate in amyloid plaques (6–10). Additionally, APP processing occurs preferentially in cholesterol-enriched domains of the plasma membrane named lipid rafts (11, 12). We previously showed that an increase of cholesterol in the plasma membrane triggers relocalization of APP-BACE1 complexes in lipid rafts and their clathrin-dependent internalization in enlarged endosomes, leading to increased APP processing and secretion of Aβ40 and Aβ42 (13–16). Cholesterol has been also described as a positive regulator of BACE1 and γ-secretase the enzyme cleaving the βC-terminal fragment (βCTF) resulting from the processing of APP by BACE1 (17, 18). Reciprocally, APP regulates cholesterol homeostasis by transcriptional regulation of key cholesterol synthesis enzyme 3-hydroxy-3-methyl-glutaryl-coenzyme A reductase (19).

Molecular simulation and physicochemical characterization showed that the Aβ5-16 segment binds to the ganglioside GM1, while the Aβ22-35 segment is linked to cholesterol in the bilayer, directing the partial insertion of the peptide in the lipid raft (20, 21). In addition, structural studies using nuclear magnetic resonance identified a cholesterol-binding site (CBS) in the transmembrane segment forming a flexible curved α-helix and in the juxtamembrane domain of the βCTF (22–24) (Fig. 1). Molecular dynamic simulations found that the APP transmembrane region and particularly the GxxxG dimerization motif was not sufficient for binding to membrane cholesterol, which also required the APP juxtamembrane segment (25). Two helical secondary structures in the βCTF fragment of APP were identified in lysophospholipid micelles (23). An R-helix that includes the transmembrane domain (TMD) extends from N698 to L723 in the sequence of APP^770^ and is terminated by three consecutive lysines. A second short R-helical segment (F690 through E693 in APP^770^ and F19 through E22 in the Aβ sequence) is located in the extracellular domain between the site of α-secretase cleavage (after K687 in APP^770^ and K16 in Aβ) and the start of the TMD. This second domain is a short reentrance loop located in a juxtamembrane region.

**Figure 1:**
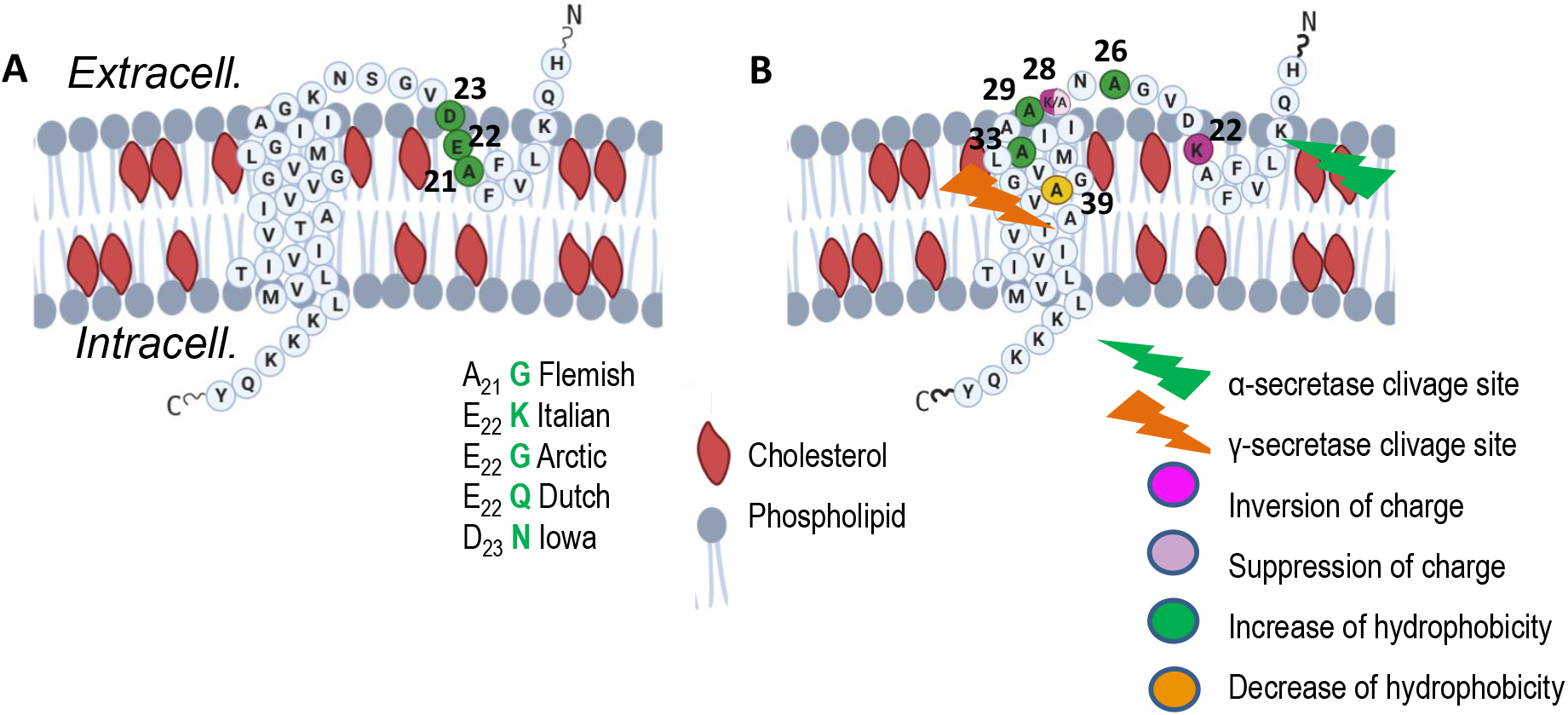
Schematic diagram of the juxta- and the trans-membrane regions of APP CTFβ within the membrane, inspired from (22). Cholesterol is highlighted in red. **(A)**: Amino acid sequence of the APP^WT^ and mutations from Familial Alzheimer Disease (FAD) cases. (**B)**: Indication of the mutations produced in the cholesterol-binding site (CBS) shown at positions 22, 26, 28, 29, 33 and 39. Numbering according to Aβ. Various colors indicate the type of changes in the mutants.

To clarify how this CBS regulates APP processing, we produced seven single mutants in the APP^751^ protein either in the juxta-membrane region at positions 22, 26 and 28 or in the TMD at positions 29, 33 and 39 of the Aβ sequence (positions 673, 677 and 679, and positions 680, 684 and 690 of APP^751^). In addition, two double mutants, 26/28 and 29/33, were constructed. We show that juxta-membrane mutants generate shorter Aβ peptides. We further demonstrate that the amount of Aβ peptides produced by one of the CBS mutants (K28A) is insensitive to the level of plasma membrane cholesterol, in distinction to APP^WT^ (13, 16). Since shorter Aβ sequences are known to be less aggregation-prone and toxic than the Aβ40 and Aβ42 peptides found in amyloid deposits (26, 27), targeting the juxta-membrane CBS-containing region of APP could be a new therapeutic option for inhibiting cholesterol-binding and hence reducing the production of toxic Aβ species.

## Methods

### Plasmid and reagents

The APP^751^ plasmid was a kind gift from Dr. Frederic Checler (IPMC, Valbonne, France). The APP-mCherry plasmid was generated by introducing the APP^751^ sequence in the pmCherry-N1 vector (Clontech) at the *XmaI/AgeI* restriction site. MβCD-cholesterol complex, bovine serum albumin (BSA), sucrose and poly-L-lysine were purchased from Sigma-Aldrich. The antibody directed against EEA1 (Early Endosome Antigen 1) was from Cell Signaling Technology. Goat anti rabbit IgG coupled to Alexa568 was from Life Technologies.

### Peptides synthesis

Amyloid peptides derived from the Aβ sequence (position 15 to 33) with an additional hydrophobic sequence (YEVH) linked to biotin were synthesized. Fmoc-amino acids, and 2-(6-chloro-1H-benzotriazole-1-yl)-1, 1, 3, 3,-tetra-methylaminium hexafluorophosphate (HCTU) were obtained from Novabiochem. N-biotine-NH(PEG)X-COOH were purchased at Merck and Sigma Aldrich. The resin and all the peptide synthesis grade reagents (N-methylpyrrolidone (NMP), N-methylmorpholine (NMM), dichloromethane, piperidine, trifluoroacetic acid (TFA), anisole, thioanisole, and triisopropylsilane) were purchased from Sigma.

Synthesis of the different Aβ Peptides were performed on a Gyros-Protein Technologies, Inc., prelude synthesizer at a 25 μmol scale using a 10-fold excess of Fmoc-amino acid relative to the preloaded Fmoc-Gly-wang-LLresin (0.33 mmol/g) or Fmoc-Ala-wang-LLresin (0.33 mmol/g). Fmoc-protected amino acids were used with the following sidechain protections: tert-butyl ester (Glu and Asp), tert-butyl ether (Ser and Tyr), trityl (His, Asn, and Gln), tert-butoxycarbonyl (Lys). Amino acids were coupled twice for 5 min using 1:1:2 amino acid/HCTU/NMM in NMP. After incorporation of each residue, the resin was acetylated for 5 min using a 50-fold excess of a mixture of acetic anhydride and NMM in NMP. Fmoc deprotection was performed twice for 3 min using 20% piperidine in NMP, and 30 sec NMP top washes were performed between deprotection and coupling and after acetylation steps. Biotinylation of APP peptide was performed on the resin after Fmoc deprotection of the N-terminal residue, using a 10-fold excess of N-biotine-NH(PEG)X-COOH and HCTU and NMM as coupling reagents (see above).

After completion, the peptidyl-resins were treated with a mixture of TFA/thioanisole/anisole/TPS/water (82:5:5:2.5:5) for 2 h. The crude peptides were obtained after precipitation and washes in cold ethyl ether followed by dissolution in 10% acetic acid and lyophilization.

Peptides were purified by reverse phase HPLC using an X-Bridge BHE C18-300-5 semi-preparative column (Waters, USA) (250 x 10 mm; 4 ml·min^−1^; solvent A, H_2_O/TFA 0.1%; solvent B, acetonitrile/TFA 0.1% using a gradient of 0-60% solvent B into A in 60 min. The purity of each peptide was checked by mass spectrometry using ESI-MS (Bruker). Lyophilized peptides were solubilized in water-acetonitrile (50%) and stored at −80 °C.

### Site-directed mutagenesis

APP-mCherry was mutated at single or double sites using the QuickChange II Site directed mutagenesis kit (Agilent) following provider-issued recommendations. Parental methylated DNA was degraded following digestion with *DpnI* and the remaining mutated DNA was transfected in competent XL1-Blue *E. Coli* (Invitrogen). Transformed Bacteria were selected on Petri dishes filled with medium containing kanamycin (30 mg/mL) (Invitrogen). In total 7 single and 2 double point mutations were produced in the cholesterol-binding site, in the juxtamembrane region of APP, at positions 22, 26, 28, 29, 33 and 39 of the Aβ sequence. Mutants are illustrated in Fig. 1B.

### Cell culture, transfection and treatments

HEK293 cells were grown in DMEM-glutamax medium (Gibco, Thermofisher Scientific) supplemented with 10% fetal calf serum (Invitrogen, CA, USA) and 1% penicillin/streptomycin (Gibco) at 37 °C and 5% CO2. Transfection experiments were carried out on HEK 293T cells with 70% confluence in OptiMEM medium (Gibco) without antibiotic using lipofectamine 2000 (Invitrogen). 0.7 μg of plasmid was diluted in 100 μL of OptiMEM medium (Gibco). In parallel, 4 μL of lipofectamine 2000 (Invitrogen) were mixed with 100 μL of OptiMEM medium. Transfection was performed according to manufacturer’s instructions (Invitrogen). After deposition of the transfection mix, the cells were placed for 4 hours in the incubator at 37 °C, washed with 1X PBS to remove any trace of transfectant and then incubated with complete medium: DMEM medium supplemented with 10% fetal calf serum and 1% penicillin-streptomycin (Gibco).

Treatment with DAPT: In order to analyze the cleavage of APP-mCherry mutants by the β- and γ-secretases, 24 hours after transfection, cells were treated for 16 hours with N-[N-(3,5-Difluorophenacetyl)-L-alanyl]-phenylglycin T-butyl ester (DAPT) (Sigma Aldrich), an inhibitor of γ-secretase, diluted to a concentration of 5 μM in complete new medium.

Cholesterol treatment: 24 hours after transfection HEK cells were washed twice with DMEM medium (Gibco), treated for 30 min with 1.4 mM MβCD-cholesterol (Sigma), dissolved in DMEM medium and then washed three times with DMEM medium.

### Protein extraction and western blot

The HEK293T cells were washed with 1X PBS and then lysed on ice using a buffer RIPA (50 mM Tris-HCl pH 8.0, 150 mM sodium chloride, 1.0% NP-40, 0.5% sodium deoxycholate, and 0.1% sodium dodecyl sulfate) (Sigma Aldrich), to which were added Phenylmethylsulfonyl fluoride 100X (Sigma Aldrich) and a cocktail of inhibitors of proteases (Complete Mini, Roche). The lysates were sonicated 3 times for 5 min then stored at −80 °C. Protein concentration of the lysates was quantified by Bradford assay (Biorad) according to the manufacturer’s instructions. Western blots were made from the cell lysates of HEK293T cells treated with DAPT. Proteins from cultured lysates HEK293T were separated in 16.5% Tris-Tricine (Biorad) polyacrylamide gels. Proteins were transferred to a polyvinylidene difluoride membrane (Biorad) at 150 V for three hours at 4 °C. After 1 hour of saturation with 10 % milk, membranes were incubated overnight at 4 °C with primary antibodies directed against APP and actin β proteins diluted to 1/2000 in BSA-azide 2 % solution. Membranes were incubated with fluorescent secondary antibodies (diluted to 1/10,000 in 0.05 % TBS-tween solution) for 1 hour under agitation at room temperature and away from direct light. The revelation and quantification of fluorescence were carried out by Odyssee’s analysis software (Set up ImageStudio CLx) (Odyssey Clx LI-COR).

### Confocal imaging

HEK293 cells cultured on poly-D-lysine-coated coverslips were fixed 24 hours after transfection using a solution of 4% paraformaldehyde in PBS for 15 min at room temperature. Cells were first incubated with anti-EEA1 antibody (1/500, Cell Signaling) and then with goat anti-rabbit secondary antibody conjugated to Alexa 488 (1/1000, Cell Signaling). Coverslips were mounted in Fluoromount medium (Southern Biotech, AL, USA). Z-stack of cells were acquired on a Fluoview FV1000 confocal microscope (Olympus, Tokyo, Japan). Fluorescence was collected with a 60x plan apochromat immersion oil objective (NA 1.35). The mean endosomes size and number per cell were analyzed with ICY software. Between 7 and 10 cells were analyzed.

### Aβ38, 40 and 42 measurements

Supernatants of HEK cells were collected on ice 24 hours after transfection in polypropylene tubes, containing phosphatase inhibitor cocktail (Complete Mini, Roche). Supernatants were then stored at −80 °C. Concentrations of the Aβ38, Aβ40 and Aβ42 species of β-amyloid peptide were measured by multiplex Electro-Chemiluminescence Immuno-Assay (ECLIA). Assays were performed according to the manufacturer’s instructions (Meso Scale Discovery (MSD) Meso QuickPlex SQ120). 100 μL of blocking buffer solution were first added to all wells to avoid non-specific binding. The plates were then sealed and incubated at room temperature on a plate shaker (450 rpm) for 1 hour. Wells were then washed three times with washing buffer, and 25 μl of the standards peptides (Aβ38, Aβ40, Aβ42) (MSD) and samples were then added to the wells, followed by an Aβ-detecting antibody at 1 μg/ml labelled with a Ruthenium (II) trisbipyridine N-hydroxysuccinimide ester (MSD). This detection antibody was 6E10 (raised against the common epitope Aβ1-16 of the human peptide, therefore the 3 Aβ38, Aβ40, Aβ42) (MSD). Plates were then aspirated and washed 3 times. MSD read buffer (containing tripropylamine as co-reactant for light generation in the electrochemiluminescence immunoassay) was added to wells before reading on the Sector Imager. A small electric current passed through a microelectrode present in each well to produce a redox reaction of the Ru^2+^ cation, emitting 620 nm red lights. The concentration of each Aβ peptide was calculated in pg/ml for each sample, using dose-response curves It was then normalized by protein concentration measured by Bradford assay (Bradford).

### Aβ (1-x) measurements

Supernatants of HEK cells were collected on ice 24 hours after transfection in polypropylene tubes, containing phosphatase inhibitor cocktail (Complete Mini, Roche). Supernatants were then stored at −80 °C. Concentrations of peptides Aβ28, 40 and 42 were measured by immune-enzymatic assay (ELISA) using the IBL Aβ(1-x) kit. This kit is a solid phase ELISA sandwich using 2 kinds of highly specific antibodies. Assays were performed according to the manufacturer’s instructions. Samples were plated in 96-well plates, containing antibodies specific for the Aβ (Precoated plate: Anti-Human Aβ(N)(82E1) Mouse IgG). Briefly, 100 μl of sample or 100 μL of Aβ40 synthetic peptides (used as the standard range, IBL) were deposited in 96-well plates. The plates were sealed and incubated overnight at 4 °C with gentle agitation. After several washings of the plates (minimum 7), 100 μl of anti-Aβ antibody solution (Mouse IgG directed against the epitope 11-28 of the human Aβ peptide, IBL) coupled to horseradish peroxidase (HRP) were deposited in the wells. The plaques were then sealed and incubated for one hour at 4 °C with gentle agitation. The plates were then washed 9 times with a wash buffer. 100 μL of a solution containing a Tetra Methyl Benzidine colorimetric agent, were deposited in the wells. After 30 minutes of incubation at room temperature and protected from light, a stop solution was deposited. Absorbance was measured at 450 nm (Thermo multiskan EX, Thermo Fisher Scientific) within 30 minutes of depositing the stop solution. The concentration of each Aβ peptide was calculated in pg/ml for each sample, using dose-response curves. It was then normalized by the protein concentration measured by Bradford assay (Bradford).

### Immunoprecipitation

Four μg of the Aβ-specific antibodies 6E10 (1-16, Biolegend) and 4G8 (17-24, Biolegend) were added separately to 25 μL of Dynabeads M-280 sheep anti-mouse (ThermoFisher Scientific) suspension, according to the manufacturer’s description. The washed antibody-bead complexes were combined (50 μL in total) and added to 3 ml supernatants of HEK cells together with 20% (v/v) Triton X-100 to a final concentration of 0.2% (m/v) and incubated overnight at +4 °C. The beads/sample complex was transferred to the KingFisher for automatic washing (in 0.2% Triton X-100, phosphate-buffered saline (PBS), pH 7.6, and 50 mM ammonium bicarbonate) and elution in 0.5% FA (v/v). The eluate was dried down in a vacuum centrifuge pending MS analysis.

### Mass spectrometry

Analysis by matrix-assisted laser desorption/ionization time-of-flight mass spectrometry (MALDI-TOF-MS) was performed using an UltraFleXtreme instrument (Bruker Daltonics) in reflector mode. Prior to analysis samples were reconstituted in 5 μl 0.1% FA/20% acetonitrile in water (v/v/v). MALDI samples were prepared using the seed layer method as previously described (28). An average of 10,000 shots was acquired for each spectrum (2000 at a time using a random walk mode). The unused sample (3 μl) was further dried down in a vacuum centrifuge and further analysed by nanoflow liquid chromatography (LC) coupled to electrospray ionization (ESI) hybrid quadrupole-orbitrap tandem MS (MS/MS), see below.

Analysis by nanoflow LC-ESI-MS/MS (Dionex Ultimate 3000 system and Q Exactive, both Thermo Fisher Scientific) was performed in a similar way as described previously (29, 30). Briefly, samples were reconstituted in 7 μl 8% FA/8% acetonitrile in water (v/v/v). An Acclaim PepMap 100 C18 trap column (20 mm × 75 μm, particle size 3 μm, pore size 100 Å, Thermo Fisher Scientific) was used for online desalting and cleanup. For separation a reversed-phase Acclaim PepMap RSLC column (150 mm × 75 μm, particle size 2 μm, pore size 100 Å, Thermo Fisher Scientific) was used. Mobile phases were 0.1% FA in water (v/v) (A) and 0.1% FA/84% acetonitrile in water (v/v/v) (B). Separation was performed at a flow rate of 300 nl/min by applying a linear gradient of 3%-40% B for 50 min at 60 °C. Spectra were acquired in positive ion mode for the mass-to-charge (m/z) range 350-1800. Both MS and MS/MS acquisitions were obtained at a resolution setting of 70,000 using 1 microscan, target values of 10^6^, and maximum injection time of 250 ms. MS/MS acquisitions were obtained using so-called higher-energy collisional dissociation fragmentation at a normalized collision energy setting of 25, an isolation window of 3 m/z units, and exclusion of singly charged ions and ions with unassigned charge.

Database search, including isotope and charge deconvolution, and peak area determination, was performed with PEAKS Studio X+ (Bioinformatics Solutions Inc.) against a custom-made APP database, which included the K28A modified sequence. All suggested fragment mass spectra were validated manually. For the label-free quantitative analysis, normalization was performed for each sample by division of the raw area with the total protein content (as determined by the Bradford assay, see above).

### Purification of exosomes from HEK293T cell culture medium

HEK293T cells were grown in DMEM 10% FCS in T150 flasks coated with Poly-lysin. At 80% confluence, cells were rinsed in PBS 1X heated at 37 °C and resuspended in OPTIMEM without FCS and incubated at 37 °C during 24 hours. Culture medium was collected in 50 mL Falcon tubes then centrifuged at 2000 g for 20 min at 4 °C. Supernatant was filtered through 0.22 μm Millipore filters and centrifuged at 100,000 g for 2 hours at 4 °C. Pellets containing the exosomes were rinsed with ice-cold PBS, centrifuged again at 100,000 g for 2 hours at 4 °C and resuspended in 75 μL ice-cold PBS. The size of exosomes was measured using Dynamic Light Scattering (mean radius 400 nm, polydispersity 25 %). Cholesterol was assessed separately using the Amplex^TM^ Red Cholesterol Assay Kit from Invitrogen (Thermo Fisher Scientific).

### Binding of amyloid peptides to exosomes

Peptides (15 μL at 20 μM) were incubated with 10 μL of exosomes (cholesterol at 30 μM) in 50 μL final volume complemented with PBS at 37 °C for 1 hour under agitation, in 2 mL Eppendorf tubes (allowing maximum agitation). Solutions were diluted in 400 μL PBS and deposited on a sucrose gradient in 5 mL tubes (from bottom to top: 3.6 mL 60 % sucrose solution in PBS, 600 μL 50 % sucrose solution and 400 μL 10% sucrose solution - the exosome + peptide mix is at the top), and centrifuged at 140,000 g for 2 hours in a swinging rotor. 11 Fractions were collected: 1 to 4 of 200 μL, 5 to 7 of 300 μL and the next 4 fractions of 1 mL. Concentrations of peptides were measured by immune-enzymatic assay (ELISA) using biotin-streptavidin-HRP detection. Assays were performed according to the manufacturer’s instructions. Briefly, 25 μl of each fraction were deposited in 96-well plates (Immunoplate, Nunc). The plates were sealed and incubated overnight at 4 °C with gentle agitation. After 3 washings of the plates in PBS, 100 μL BSA 1% in PBS was added for one hour followed by 3 washings in PBS. Streptavidin solution coupled to horseradish peroxidase (HRP) (Sigma) in PBS (1/2500 dilution) were deposited in the wells (100 μL). The plaques were then sealed and incubated for one hour at 4 °C with gentle agitation. The plates were then washed 3 times with 0.05 % PBS-tween solution. After 30 minutes of incubation at room temperature with 3,3’, 5, 5’-tetramethylbenzidine (TMB Peroxidase EIA Substrate Kit, Biorad), a stop solution (H2SO4 2M) was deposited. Absorbance was measured at 450 nm (Varioscan) within 30 minutes of the deposition of the stop solution.

### Statistical analysis

All analyses were performed using GraphPad Prism version 6.00 for Windows. Statistical tests were two-tailed and conducted at a 5% significance level. Student t test was performed to compare mutant vs control condition and non-parametric Mann and Whitney t test was applied for small datasets (n≤4).

Data were derived from 3 independent cultures with 2 to 3 replicates per culture, except confocal imaging which was performed on 1 culture and 7 to 10 cells per condition. Data are presented as mean ± SEM.

## Results

### Production of APP mutants of the cholesterol-binding site (CBS)

Starting from the structural study that characterized the CBS on APP (22) (Fig. 1A), we used site-directed mutagenesis to produce point mutations in the APP^751^mCherry plasmid. In total seven mutants were generated in the juxtamembrane region at positions 22, 26 and 28, and in the transmembrane region at positions 29, 33 and 39 of the Aβ sequence (positions 673, 677 and 679, and positions 680, 684 and 690 of APP^751^). In addition, we produced two double mutants at juxtamembrane positions 26/28 and at transmembrane positions 29/33 (Fig. 1B). Point mutations corresponded to either inversion of charge of the mutated amino acid such as in the E22K and the K28E mutants, suppression of charge as in the K28A mutant, decrease of hydrophobicity as in the V39A mutant or increase of hydrophobicity as in the S26A, G29A and G33A mutants (Fig. 1B). These changes of either charge and/or hydrophobicity at these positions have been shown by NMR to alter the cholesterol-binding properties (22). All plasmids were sequenced and the mutations confirmed.

### Dosage of Aβ peptides produced by HEK293T cells transfected with APP mutants of the CBS

We used the Meso Scale Discovery multiplex ELISA for dosage of Aβ38, Aβ40 and Aβ42 using capture antibodies recognizing the C-terminal part of the Aβ sequence (neoepitope ending at amino acid 38, 40, 42, respectively) and a detection antibody binding to the N-terminal part of Aβ (4–9) by Electro-Chemiluminescence. All mutations produced were localized outside the epitopes of both the capture and the detection antibodies. As described previously, we found that the levels of Aβ40 secreted by HEK293 cells were higher than the levels of secreted Aβ42 (data not shown) as suggested earlier (18, 31). We found that mutations at positions 28 and 39 dramatically reduced the secretion of Aβ40 and Aβ42 as compared with APP^WT^, while the effect was weaker with mutations at positions 22, 26, 29 and 33 (Fig. 2A,B). Secretion of Aβ38 was undetectable with all transfected plasmids. We then asked whether the large decrease in Aβ40/42 secretion by HEK293T cells transfected with mutated APP^751^ was due to an accumulation of intracellular peptides or to differences in APP processing. Since the MSD Multiplex assay can only quantify full length Aβ38, Aβ40 and Aβ42, we used the IBL ELISA with a capture antibody that targets the N terminus of Aβ peptides and a detection antibody raised against Aβ11-28 of Aβ, able to measure shorter Aβ peptides (Aβ1-x with x≥16) in addition to Aβ38, Aβ40 and Aβ42. Using this IBL kit, we found that mutants at positions 26 and 28 and the double mutant 26/28 secreted significantly more Aβ1-x peptides with x≥16 while all other mutants showed decreased secretion (Fig. 2C). As the secretion of Aβ40 and Aβ42 was reduced using these mutants (Fig 2A, B), we concluded that mutants at positions 26 and 28 and the double mutant 26/28 secreted shorter peptides within the range Aβx-16 to Aβx-42.

**Figure 2:**
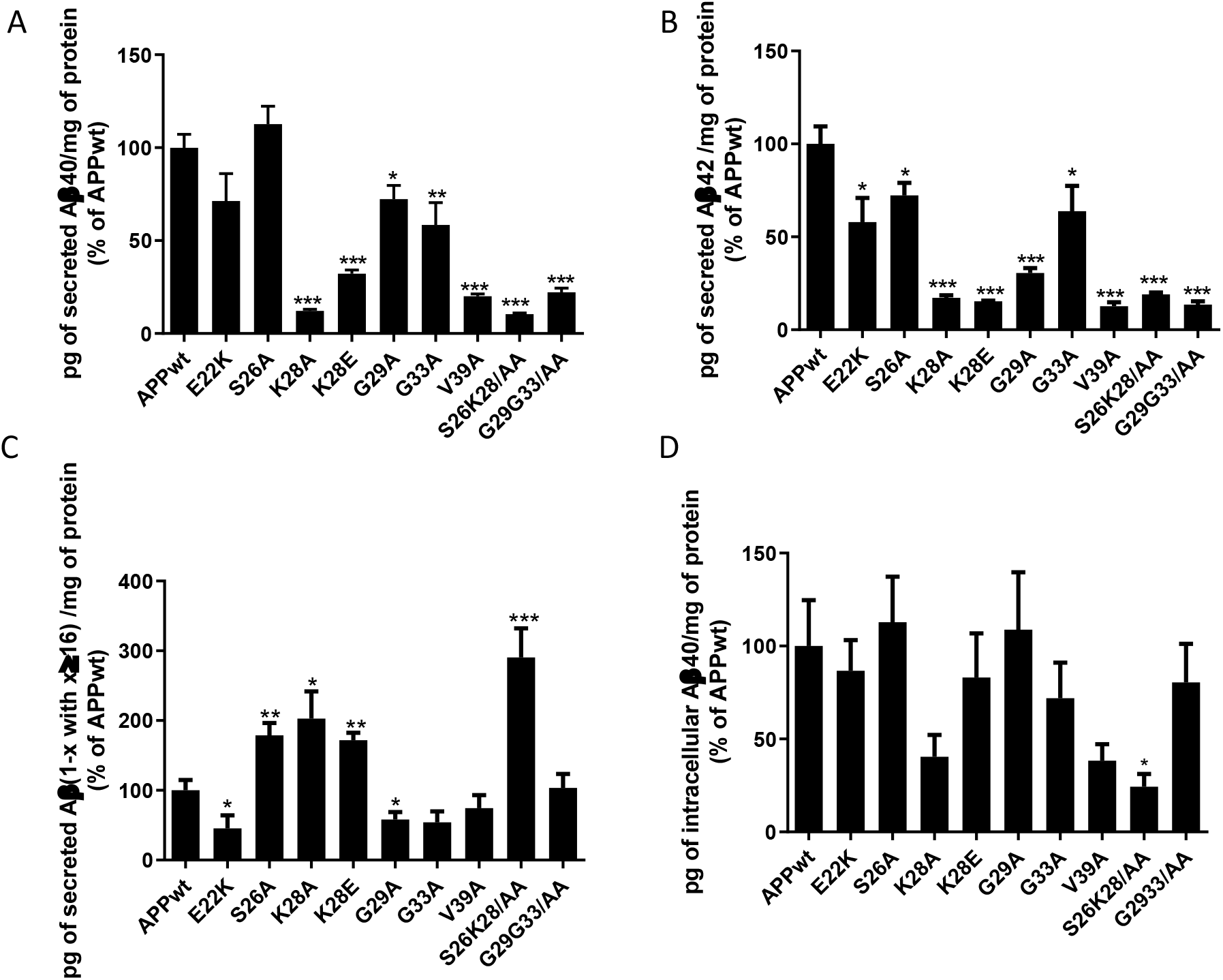
Mutations in the CBS modulate the Aβ secretion in transfected HEK293T cells without accumulation in the intracellular compartments. (**A**, **B, D)**: MSD assay of extracellular Aβ40 and Aβ42 levels respectively, on transfected HEK293T with different mutants of the APP CBS. The results are normalized with the amount of intracellular proteins determined by Bradford assay and represented as a percentage of Aβ produced by APP^wt^. (**C)**: ELISA (IBL kit) for Aβ(1-x with x≥16) extracellular levels on HEK293T transfected with the different APP mutants. The results are normalized with the amount of intracellular protein determined by Bradford assay and normalized to percentage of Aβ produced by transfected HEK293T cells with APP^wt^. Numbering according to Aβ (t test, mean +/− SEM, *: p <0.05, **: p <0.01, ***: p <0.001, comparison with APP^wt^, 3 independent cultures with 2 to 3 replicates per culture 6 <n <9).

Altogether, we showed that mutating the CBS of APP at positions 26 and 28 of the Aβ sequence altered the size of Aβ peptides produced while mutations at positions 22, 29, 33 and 39 decreased the amount of secreted Aβ peptides ranging from Aβ16 to Aβ42. Mutation at position 28 gave the strongest effect compared to mutation at position 26, while the double mutant (positions 26 and 28) showed cumulative effect, particularly for shorter Aβ peptides (Fig. 2C).

We then asked whether the large decrease in Aβ40 and Aβ42 secretion was due to an accumulation of intracellular peptides or to differences in APP. We used the MSD assay to measure intracellular Aβ38, 40 and 42. In Fig. 2D, we found again less although non-significant accumulation of the intracellular Aβ40 in HEK293T cells transfected with the mutants at positions 28 and the double mutant 26/28 than with the APP^WT^. Levels of Aβ42 and Aβ38 were undetectable.

### Processing of APP^WT^ and APP^K28A^

We then characterized the APP mutant K28A (APP^K28A^) which showed the most contrasted profile of Aβ secretion. Because of the large decrease of Aβ40 and Aβ42 secretion observed in HEK293T cells transfected with APP^K28A^, we tested the activity of the α-, β-, and γ-secretases by quantifying the levels of full length APP, C-terminal fragments (βCTF) and AICD (APP intracellular domain) in the presence or absence of the γ-secretase inhibitor, the dipeptide DAPT. In the absence of DAPT the activities of α-and β-secretases are measured through the production of AICD (Fig. 3C). In the presence of DAPT, production of AICD is inhibited, thus allowing quantification of βCTFs reflecting the activity of β-secretase only (Fig. 3B). Using western blot quantification, Fig. 3 shows that the levels of full length APPmCherry did not vary between transfections (data not shown). Moreover, βCTFs were significantly but modestly decreased in cells transfected with APP^K28A^ compared to cells transfected with APP^WT^ while the levels of AICD remained unchanged. Of note, the small decrease of βCTFs observed with APP^K28A^ in Fig. 3 did not compare with the dramatic drop in secretion of Aβ40 and 42 (Fig. 2A-B) or in intracellular expression of Aβ40 (Fig. 2D) within cells transfected with APP^K28A^ compared to cells transfected with APP^WT^. Overall, α-, β-, and γ-secretases activities were not modified by CBS mutation.

**Figure 3:**
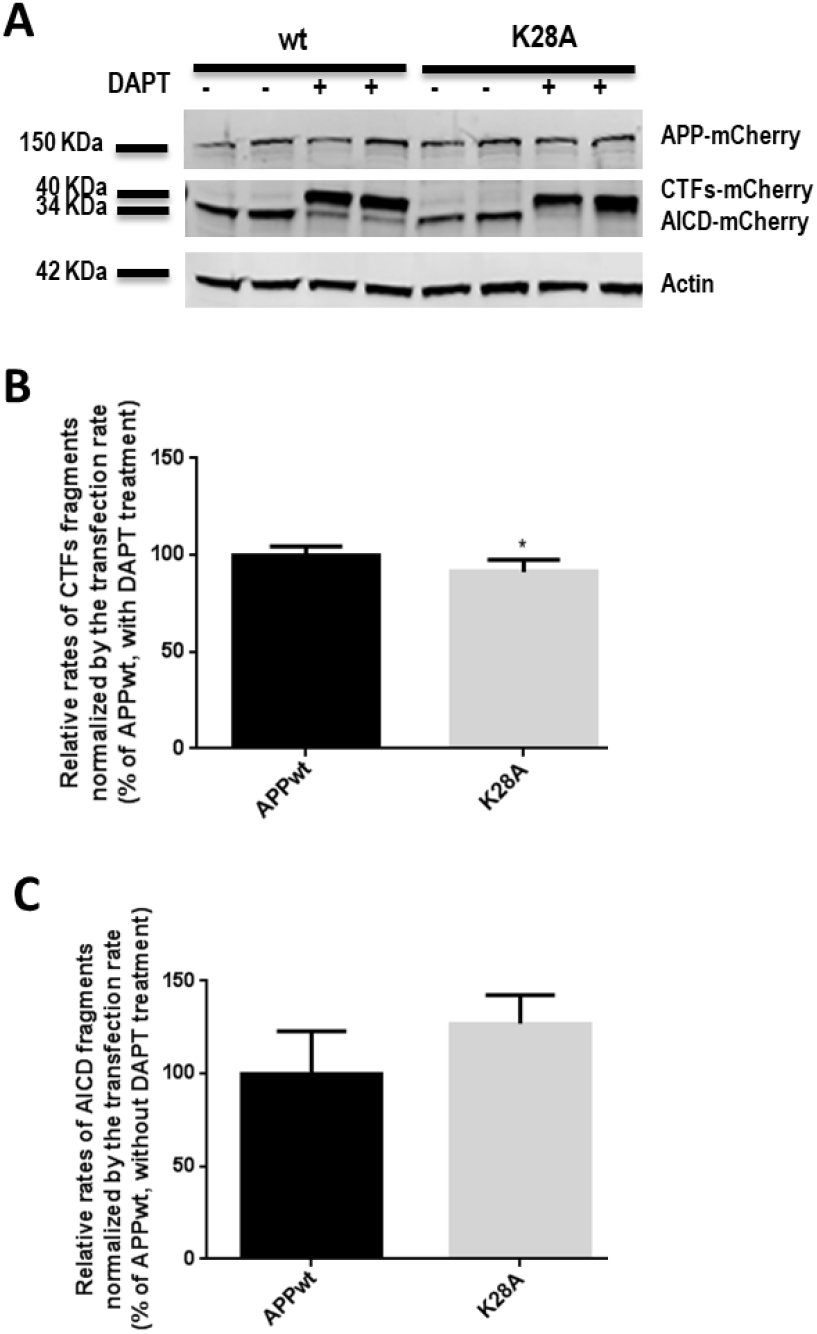
Effect of K28A mutation in the CBS on the activity of α, β and γ-secretase in transfected HEK293T. (**A)**: Western blot of HEK cell lysates transfected with APP^wt^ and APP^K28A^. The relative levels of CTFs (**B**) and AICD (**C**) fragments generated by the juxtamembrane mutation K28A are normalized by transfection rates (level of APP-mCherry). (Test Mann Whitney, mean +/− SEM, *: p = 0.0260, 3 independent cultures with 4 <n <6).

### Subcellular localization of APP^WT^ and APP^K28A^

Since APP processing occurs in the endolysosomal compartment, we compared the percentage of colocalization of APP^WT^-mCherry and APP^K28A^-mCherry with an anti-EarlyEndosomeAntigen1 (EEA1) antibody in transfected HEK293T cells. Fig. 4 clearly shows that APP^WT^ and APP^K28A^ similarly localized in the early endosomal compartment, strongly suggesting that subcellular localization of APP^WT^ and APP^K28A^ in early endosomes was comparable.

**Figure 4:**
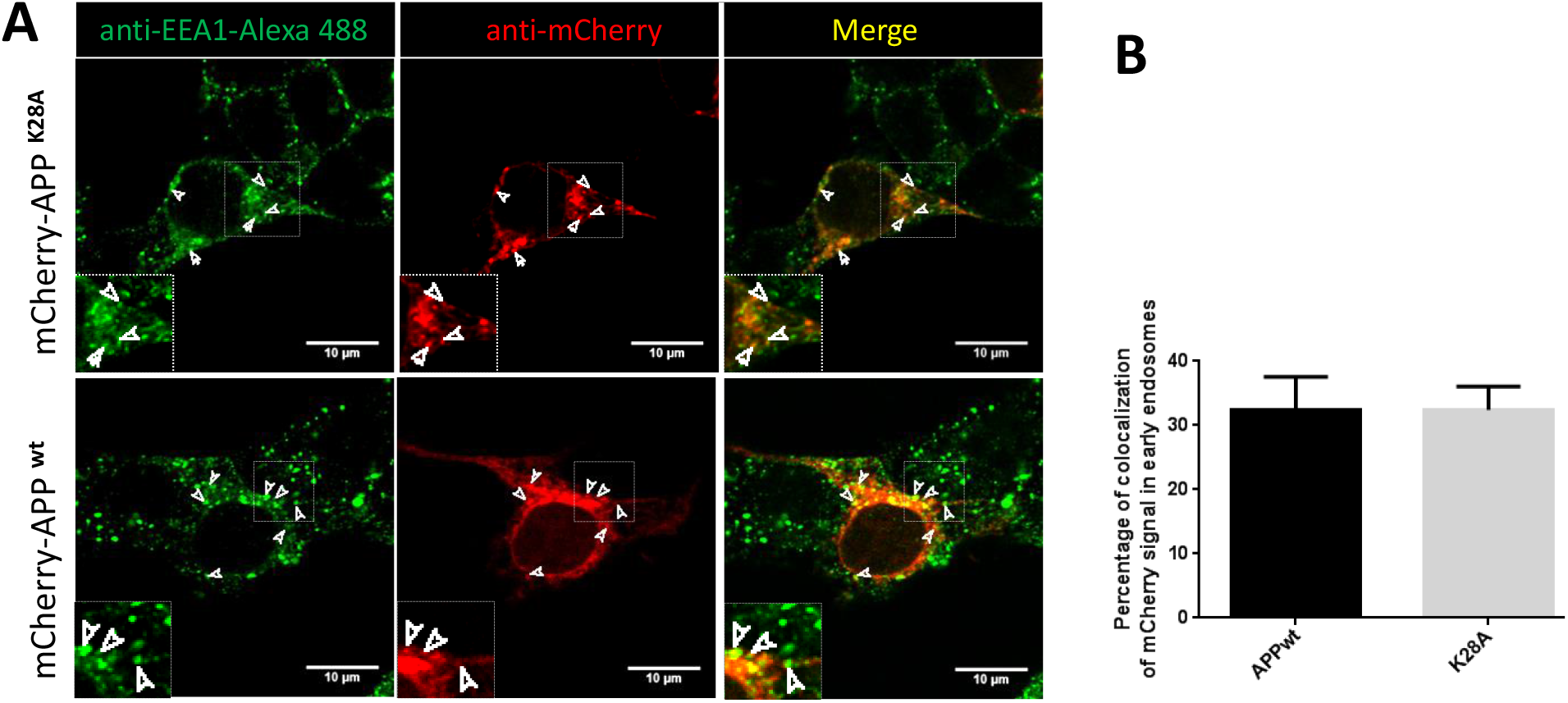
Effect of K28A mutation on endosomal localization of APP-mCherry. Effect of K28A mutation on endosomal localization of APP-mCherry. HEK293T cells were transfected with APP^wt^-mCherry or APP^K28A^-mCherry, fixed after 24 hours and immunolabelled for early endosomes using an anti-EEA1 antibody. (**A**) Arrows indicate APP^wt^-mCherry or APP^K28A^-mCherry in EEA1-positive early endosome. **(B)** Percentage of colocalization of the two fluorescent signals mCherry and Alexa488 was quantified from single stack of images acquired on the confocal microscope. The values are given as mean ± SEM **(B)**. Statistical differences were analyzed by t test, 7 <n <10 cells/experiment.

### Effect of cholesterol treatment of stable HEK293T clones expressing APP^WT^ and APP^K28A^ on Aβ40 and Aβ42 production

We wanted to test whether the addition of cholesterol in the plasma membrane, known to trigger APP processing and Aβ production, was able to do so in cells expressing the APP^K28A^ mutant. We first produced HEK293T clones stably expressing APP^WT^ and APP^K28A^ and selected those which expressed similar levels of the protein APP-mCherry (clones 1H4 and 2E5 in Supplementary Fig. 1). We confirmed that clone 2E5 expressing APP^K28A^ produced significantly less Aβ40 and Aβ42 peptides as compared to clone IH4 that expressed APP^WT^ (Fig. 5 and Supplementary Fig. 1). HEK293T clones stably expressing APP^WT^ and APP^K28A^ were treated with methyl-β-cyclodextrin (MBCD) loaded with cholesterol as described previously thus inducing a 15-20 % increase of membrane cholesterol concentration (14, 16). While cholesterol increase at the plasma membrane raised Aβ40 and Aβ42 secretion in APP^WT^ expressing clone, APP^K28A^ clone expressing the CBS mutant, as expected, was insensitive to cholesterol changes (Fig. 5).

**Figure 5:**
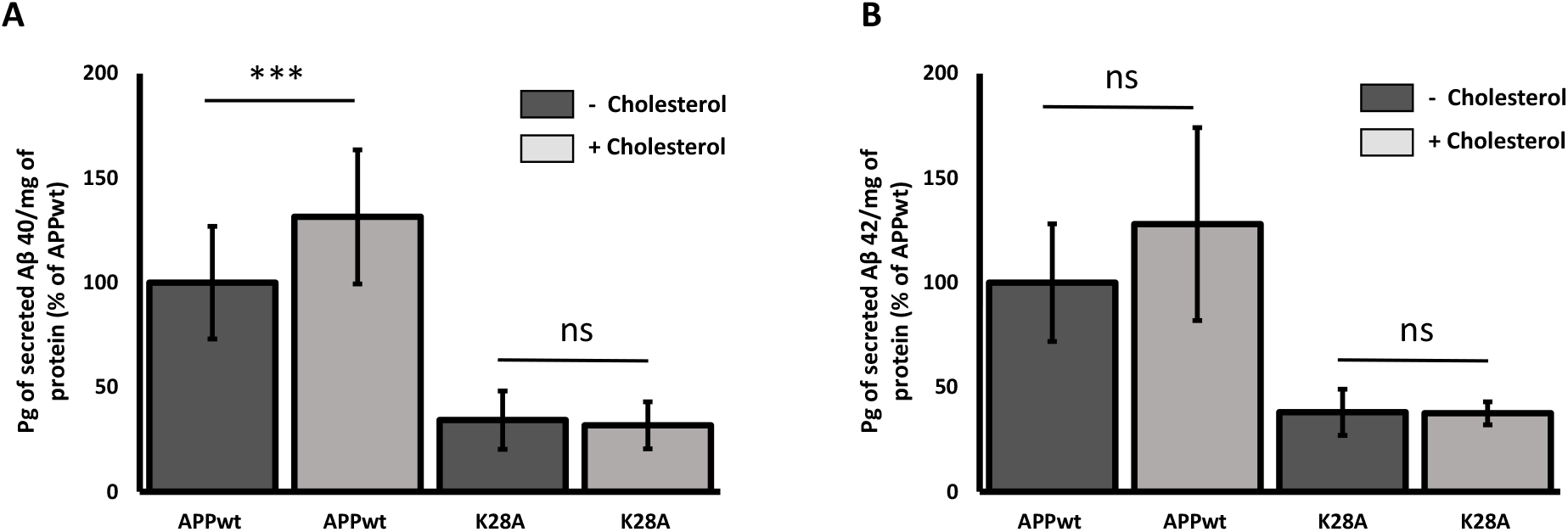
The K28A mutation in the CBS of APP abolishes the modulation of Aβ secretion by cholesterol increase at the plasma membrane. MSD assays for secreted Aβ40 **(A)** and Aβ42 **(B)** levels from HEK293T clones stably expressing APP^WT^ and APP^K28A^ in the absence and in the presence of additional methyl-β-cyclodextrin (MBCD) loaded with cholesterol. The results are normalized with the amount of intracellular proteins determined by Bradford assay and represented as a percentage of Aβ produced by APP^WT^ in the absence of MBCD loaded with cholesterol. (Test two-way ANOVA, 4 independent experiments/culture with 2 <n <6).

### Effect of cholesterol treatment of stable HEK293T clones expressing APP^WT^ and APP^K28A^ on Aβ peptides profile using mass spectrometry

In order to confirm that K28A mutation in APP changes the Aβ profile of peptides secreted from longer towards shorter size, we analyzed the supernatants of HEK293T clones stably expressing APP^WT^ and APP^K28A^ using MALDI-TOF-MS and LC-ESI-MS and - MS/MS. A different Aβ peptide pattern was observed between APP^WT^ and APP^K28A^ samples (Figure 6). In APP^WT^ samples the full length Aβ1-40 is present, together with several N-truncations of Aβx-37/38/39/40, while these peptides are absent in the APP^K28A^ samples, where Aβx-33/34 are the major forms identified. A full list of the peptides identified by LC-ESI-MS/MS is shown in Supplementary Table. Cholesterol treatment of HEK293 cells transfected with APP^WT^ or APP^K28A^ did not modify the profile of secreted Aβ peptides.

**Figure 6:**
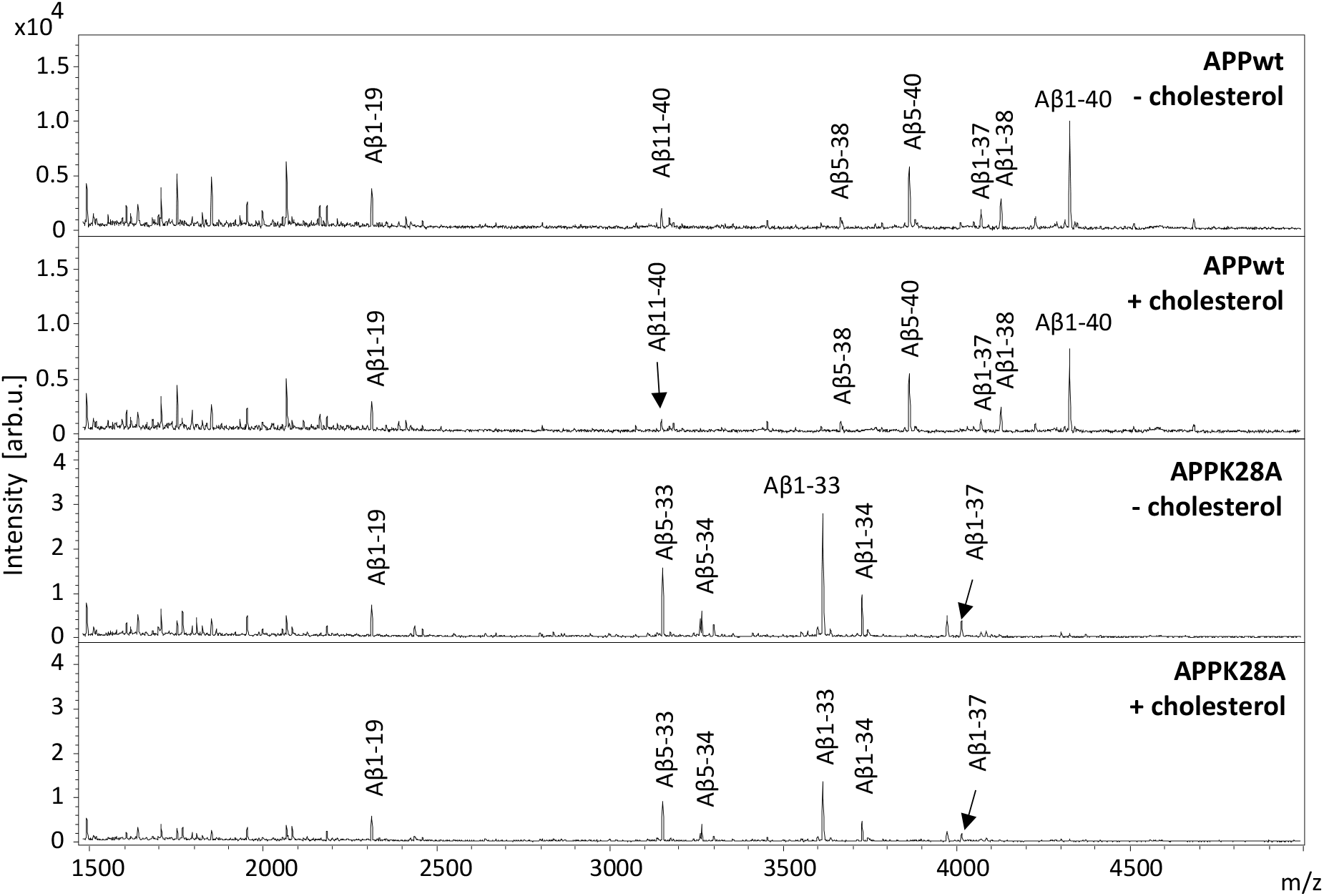
MALDI mass spectra of APP^WT^ and APP^K28A^ both treated with cholesterol (+cholesterol) and untreated (-cholesterol). An Aβ pattern difference between APP^WT^ and APP^K28A^ is observed, with AβX-37/38/39/40 present in WT while AβX-33/34 peptides are the most abundant in APP^K28A^.

### Binding of Aβ-derived peptides to membranes of exosomes purified from HEK293T cells

Next we wanted to test whether mutations of the CBS in the juxtamembrane segment of APP could change the interaction with lipid bilayers formed by natural membranes. Three peptides corresponding to the juxtamembrane region of APP (positions 15 to 33 on Aβ) were synthetized with a linker and a biotin (Fig. 7). In addition to Aβ15-33^WT^ and Aβ15-33^K28A^, we synthetized Aβ15-33^E22K^ peptide carrying the Italian mutation found in familial cases of AD. Cholesterol-rich exosomes secreted by untransfected HEK293T cells were purified by ultracentrifugation and incubated with the three peptides for 1 hour at 37 °C. Peptides bound to exosomes were separated on a sucrose gradient and quantified by luminescence with streptavidin coupled to HRP. Fig. 6 shows that Aβ15-33^WT^ bound specifically to exosomes while Aβ15-33^E22K^ and Aβ15-33^K28A^ did not, thus confirming that K28A mutation in the CBS alters the binding of Aβ15-33 peptides to cholesterol containing natural membranes.

**Figure 7:**
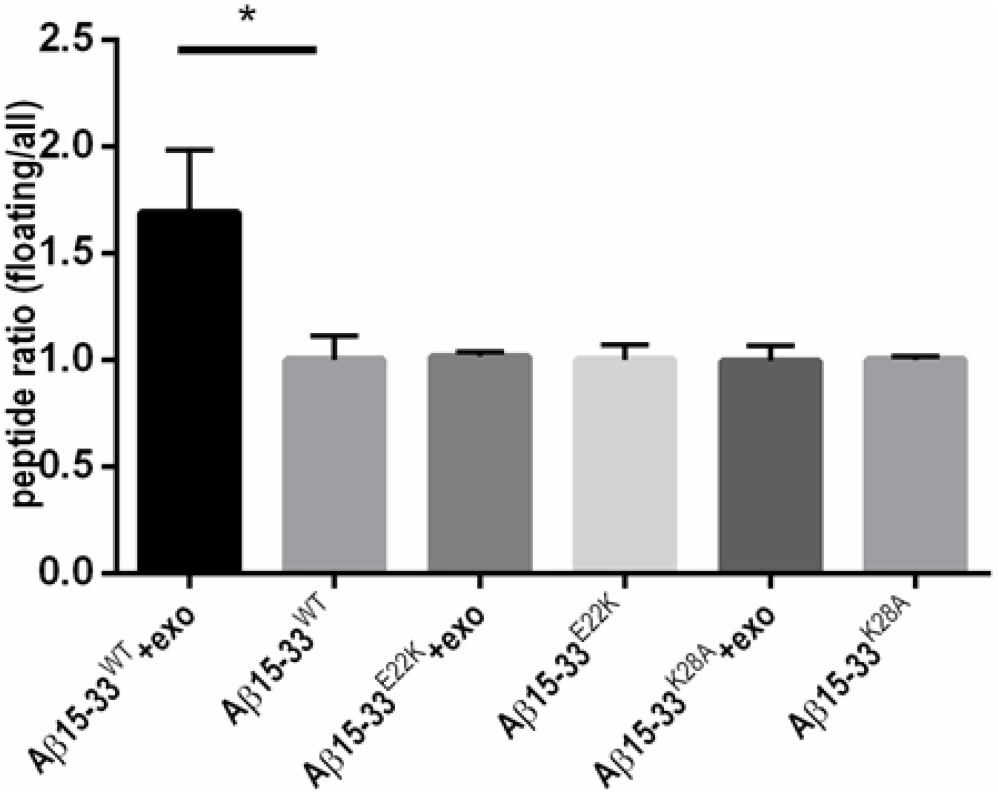
Effect of mutations of the CBS in the juxtamembrane segment of APP on their binding to exosomes. Evaluation of the amount of biotinylated peptides floating in an 11 mL sucrose gradient after incubation with and without exosomes and ultracentrifugation. Biotinylated peptides were quantified by ELISA assay using HRP-streptavidin (n=3). Ratio of bound peptides corresponds to biotine signal in the first 4 fractions (0 and 10% sucrose) divided by the total signal of the 11 fractions (0 - 10 - 50 and 60% sucrose) (mean +/− SD, * p<0.05, two-tailed P value n=3).

## Discussion

Our previous work demonstrated that membrane cholesterol is an important factor modulating the processing of the transmembrane protein APP and hence the production of toxic Aβ peptides. We found that transient increase of cholesterol at the plasma membrane of non-neuronal and neuronal cells induces a rapid relocalization in lipid rafts of APP and the β-secretase BACE1, the first APP-processing enzyme of the amyloid pathway (14, 15). This was followed by a rapid internalization of APP-BACE1 complex in endosomes with larger size and an increase of Aβ40/42 secretion (14–16).

We show here that mutations in the transmembrane or in the juxtamembrane regions of APP involved in cholesterol-binding differentially regulate APP processing. Point mutations in the APP transmembrane domain modulating the hydrophobicity of specific residues at position 29 (G29A), 33 (G33A) involved in the GxxxG dimerization motifs, and 39 (V39A) of the Aβ sequence lead to a significant reduction of the secretion of Aβ peptides (Aβ40 and Aβ42 as well as shorter Aβ1-x with x≥16 peptides, which we show was not due to any accumulation of intracellular Aβ peptides. In addition, the double mutation G29A and G33A provided more dramatic and cumulative effects on Aβ secretion. Reduction of Aβ secretion by mutations in the glycines of the GxxxG could thus be due not only to differences in cholesterol binding but also result from inhibition of the γ-secretase as was found previously (32). Interestingly, here we show that point mutations in the APP juxtamembrane domain producing either an increase of hydrophobicity (S26A) or a change of charge (E22K, S26A, K28A and K28E) had different effects on Aβ secretion. While the E22K mutant produced significantly less Aβ42 and Aβ28-42, all three others mutants (S26A, K28A and K28E) showed lower levels of Aβ42 but higher levels of Aβ1-x with x≥16 peptides, with K28A giving the most contrasted differences. Interestingly the secreted Aβ peptide profile from the double mutant S26A and K28A was more similar to K28A profile as compared to S26A thus indicating that the K28A mutation prevailed.

Two important residues from the juxtamembrane segment namely E22 and D23 have been found to control pH-dependant binding of cholesterol to APP (33). At low pH, such as in the endosomal compartment, E22 and D23 are neutral and bind cholesterol thus allowing APP processing by the β- and γ-secretases while change of charge as in the E22K mutant alters this processing. This glutamic acid at position 22 of Aβ is mutated in several dominant familial cases of AD (FAD) surprisingly showing low levels of Aβ peptides. As mirrored here in cellular models, the Italian mutation E22K produces low levels of Aβ42 in the brain of affected individuals with prominent cerebral amyloid angiopathy and lack of neurofibrillary tau and neuritic plaques (34, 35). Here we found that binding of the corresponding synthetic mutated peptides derived from the juxtamembrane region of Aβ (Aβ15-33^E22K^) to lipid bilayers formed by cholesterol rich natural membranes was largely decreased as compared to Aβ15-33^WT^ peptide. This change of interaction could induce the juxtamembrane loop to exit the plasma membrane. Indeed, it was shown previously that the Aβ22-35 region is linked to cholesterol in the lipid bilayer (36). In Fig. 1 we represent equal levels of membrane cholesterol in the two leaflets. However, the exact ratio of cholesterol between outer and inner leaflets has not been definitively clarified (37) but several authors suggest that there could be ten times more cholesterol in the outer leaflet than in the inner leaflet (38, 39). This later property could favour the binding of the APP juxtamembrane region to the enriched cholesterol leaflet.

Other pathological mutations at position E22 have been identified and show changes in Aβ formation. Arctic mutation carriers (E22G) have lower levels of Aβ40/42 while Aβ protofibrils are increased (40). In addition, individuals carrying the E22Q Dutch mutation show hereditary cerebral haemorrhage with Dutch-type amyloidosis with characteristic cerebral amyloid angiopathy and an increased formation of oligomeric and fibrillar Aβ (35). Change of binding of cholesterol following mutations at position E22 remains to be demonstrated in vivo.

The profile of Aβ secreted by S26A, K28A and K28E mutants of APP were similar to each other with a decrease of Aβ42 and an increase of Aβ28-42 suggesting that shorter Aβ peptides are produced by these mutants. In addition, other mutants at positions S26 and K28 namely S26L and K28S were also found to produce large decrease of Aβ secretion (41). Using MS methods we confirmed that the K28A mutant produced mostly Aβ33 and Aβ34 peptides with no detectable levels of Aβ40 or Aβ42, in line with a previous study (42). These authors suggested that the change in the length of secreted Aβ peptides induced by the K28A mutation was due to a shift in the primary cleavage site of the γ-secretase without significant change to the ε cleavage occurring between L49 and V50 and producing AICD. In this model, the lysine at position 28 anchors the Aβ sequence at the juxtamembrane thus limiting the accessibility to the stationary transmembrane γ secretase. Mutating this lysine would suppress the anchoring thus allowing entry in the transmembrane γ secretase active site (42).

Here we propose that interaction of lysine at position 28 with membrane cholesterol participates in the anchoring of the Aβ22-28 sequence at the juxtamembrane. Mutating this lysine at position 28 suppressing cholesterol-binding will “de-staple” the Aβ region of APP, thus allowing the entry of the sequence in the membrane and permitting cleavage by γ-secretase at positions 33-34. Previous molecular dynamics simulations have shown that K28A mutation reduces intrapeptide hydrophobic interactions between E22/D23 and K28 (43) and increases helix content while decreasing β-sheet and hydrophobic contacts and electrostatic interactions (44, 45). The K28A mutation would thus not only reduce the interaction with cholesterol but also decrease intrapeptide interactions. Evidence for this may be summarized as follows.

First, we show that APP^K28A^ produces shorter size Aβ peptides (Aβx-33/34) without any change in the levels of AICD. Second, we found that the APP^K28A^ mutant is insensitive to the effect of membrane cholesterol increase on Aβ secretion with no change in the levels nor the profile of Aβ peptides. According to our previous work (14–16) and the data presented here, transient membrane cholesterol increase triggers APP^WT^ processing and Aβ40/42 secretion. Remarkably, this effect of cholesterol was not observed with the APP^K28A^ mutant. This change was unlikely due to difference of endocytosis since the subcellular localization of APP^K28A^ and APP^WT^ in the endosomal compartment were similar. Third, binding of the corresponding synthetic mutated engineered peptides derived from the juxtamembrane region of Aβ (Aβ15-33^K28A^) to lipid bilayers formed by natural membranes was largely decreased as compared to Aβ15-33^WT^ peptide. Thus the affinity of Aβ15-33^K28A^ and Aβ15-33^E22K^ for the cholesterol rich membrane is reduced, therefore their tilting within the membrane and in particular within the outer leaflet should be modified. Consequently, these CBS mutants appear to different positions in the plasma and intracellular membranes and induced changes in their cleavage by γ-secretase produces shorter Aβ peptides.

Aβ34 is produced through degradation of Aβ40 and 42 but not βCTF by BACE1 (46, 47). It is present in the brain of AD patients and 3xTg mice (48) and is elevated in individuals with mild cognitive impairments at risk for dementia, correlating with amyloid positivity and pericyte mediated clearance (49–51). Here we show that APP cleavage by BACE1 is not affected by the K28A mutation. However, whether degradation of Aβ40^K28A^ and Aβ42^K28A^ by BACE1 is favoured remains to be shown using synthetic peptides and reconstituted BACE1 protein. However it is quite unlikely since we show that K28A mutation mostly produces Aβ33 peptides that are not degraded from Aβ34 by the matrix metalloproteases MMP-2 and MMP-9 (52).

## Conclusions

Collectively, the present data demonstrate that specific mutations in the juxtamembrane segment of the cholesterol-binding site of APP lead to the production of shorter Aβ peptides, likely by altering the anchoring and positioning of APP in the plasma membrane. By allowing access to the active transmembrane γ secretase site, this results in an increase in APP processing. A tight regulation of membrane cholesterol is particularly important at the presynaptic terminals of neuronal cells where APP is enriched and its expression, distribution and processing are regulated by synaptic activity (53, 54). Thus, together with neuronal activity, the cholesterol content of membranes and the CBS on APP could play a critical role in regulating APP processing and the resulting amount, size and toxicity of Aβ peptides.

These observations underpin interest in central cholesterol as a potential target for the treatment of AD and several strategies have proven successful in preclinical models (55). For example, CYP46A1 overexpression to accelerate brain cholesterol clearance reduces amyloid pathology and improves cognitive deficits in murine models for AD (56). Intriguingly, and suggesting the broader relevance of cholesterol, in cultured neurons cholesterol has been shown not only to regulate Aβ secretion through its interaction with the APP CBS, but also tau pathology *via* a different mechanism involving the proteasome (57). The present results establish a role of the juxtamembrane region of APP containing the CBS as a regulator of the size of Aβ peptides produced by processing of APP processing: this specific region could then be a novel target for inhibiting cholesterol-binding and reducing the production of toxic Aβ species. K28 has been targeted using lysine-specific molecular tweezers to inhibit assembly and toxicity of amyloid peptides (58–60). It will be of interest to determine whether editing of this residue using CRISPR Cas9, antisense knockdown of the mutant protein, or treatment with antibodies targeting the juxta-membrane region will alter APP processing.

## List of abbreviations

Aβ: amyloid-β
AD: Alzheimer’s disease
AICD: APP intracellular domain
APOE: apolipoprotein E
APP: Amyloid Precursor Protein
CBS: cholesterol-binding site
βCTF: C-terminal fragment
DAPT: *tert*-Butyl (2*S*)-2-[[(2*S*)-2-[[2-(3,5-difluorophenyl)acetyl]amino]propanoyl]amino]-2-phenylacetate
EEA1: EarlyEndosomeAntigen1
MBCD: methyl-β-cyclodextrin
TMD: transmembrane domain

## Declaration

The datasets generated and/or analysed during the current study are available from the corresponding author on request.

## Authors’ contribution

LB, MCP, CM, BS, NG and LH made substantial contributions to the conception and design of the work; LH, BS, AK, GF, EG, GB, participated in the acquisition, analysis, and interpretation of data; MCP, LB, BS, EP, KB, HZ, MJM have drafted the work or substantively revised it and all authors have approved the submitted version and have agreed both to be personally accountable for the author’s own contributions and to ensure that questions related to the accuracy or integrity of any part of the work, even ones in which the author was not personally involved, are appropriately investigated, resolved, and the resolution documented in the literature.

## Acknowledgements

The authors wish to thank Inger Lauritzen and Frédéric Checler from IBPC Sophia Antipolis for protocols for exosome purification and important discussions on Aβ34 respectively, Karen Perronet and Julien Moreau from Institut d’Optique Paris-Saclay for useful discussions and Bernadette Allinquant and Serge Marty for critical reading of the manuscript. This work was supported by ‘Investissements d’avenir’ ANR-10-IAIHU-06, Institut de Recherche Servier. LH was supported by a fellowship from Institut de Recherche Servier. HZ is a Wallenberg Scholar supported by grants from the Swedish Research Council (#2018-02532), the European Research Council (#681712), Swedish State Support for Clinical Research (#ALFGBG-720931), the Alzheimer Drug Discovery Foundation (ADDF), USA (#201809-2016862), and the UK Dementia Research Institute at UCL.

---

**Supplementary Figure:**
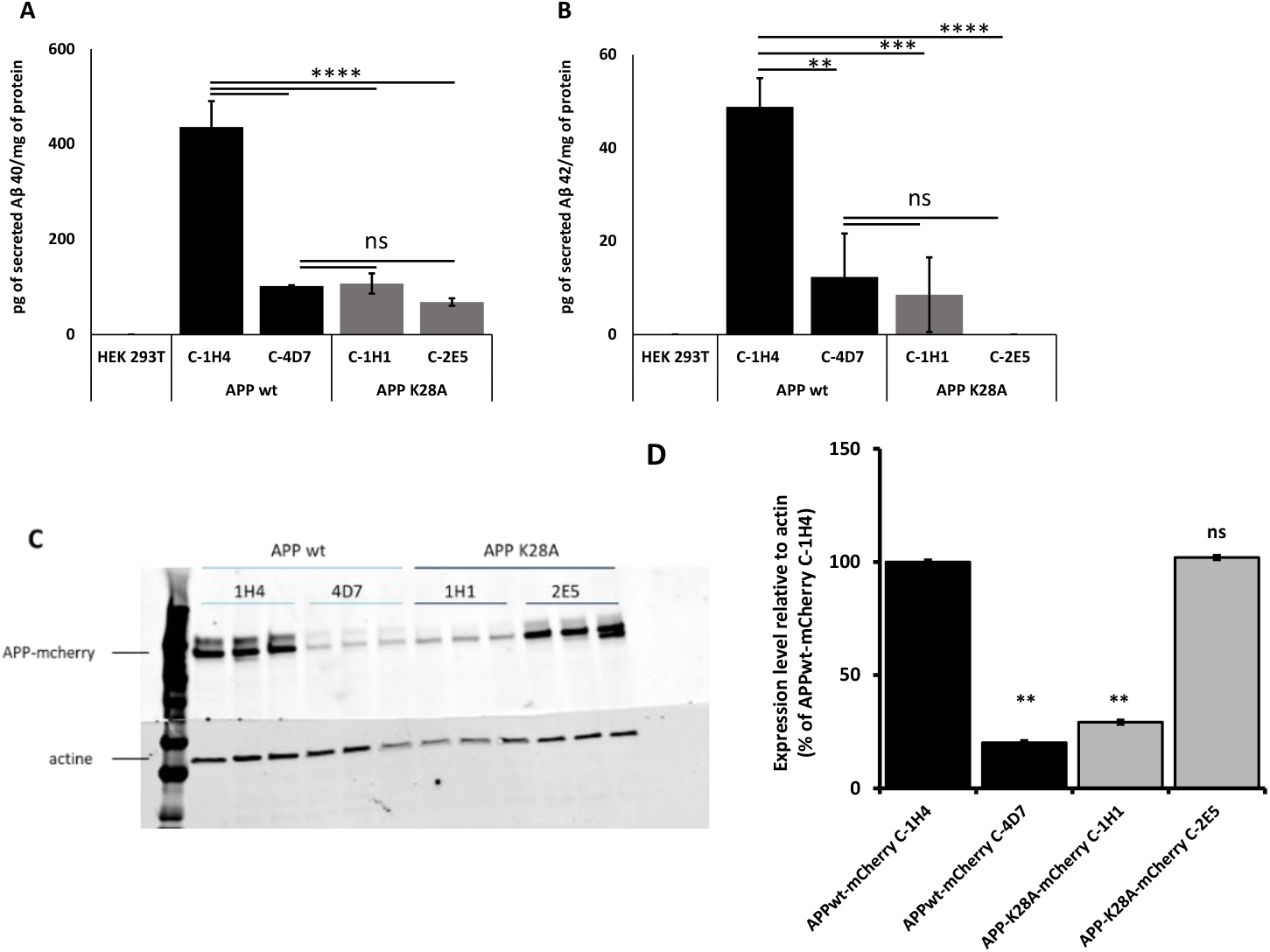
Characterization of HEK293T clones stably expressing APP^WT^mCherry and APP^K28A^mCherry. **A**, **B**: MSD assay for secreted Aβ40 and Aβ42 respectively. No Aβ40 and 42 were detected in the HEK293T untransfected control clones. One-way ANOVA Tukey’s multiple comparisons (n=3, mean ± standard error); **C**: Western blot analysis of mcherry expression levels of clones stably expressing APPwt-mCherry and APPK28A-mCherry were detected with an anti-mCherry antibody. D: Expression levels of APPmCherry were quantified using ImageJ. Expression levels relative to actin were normalized to the APPmCherry level of each clones. Statistics: two-way ANOVA (n=3, mean ± standard error). ****p<0.0001***p<0.001** p<0.01

**Supplementary Table.**
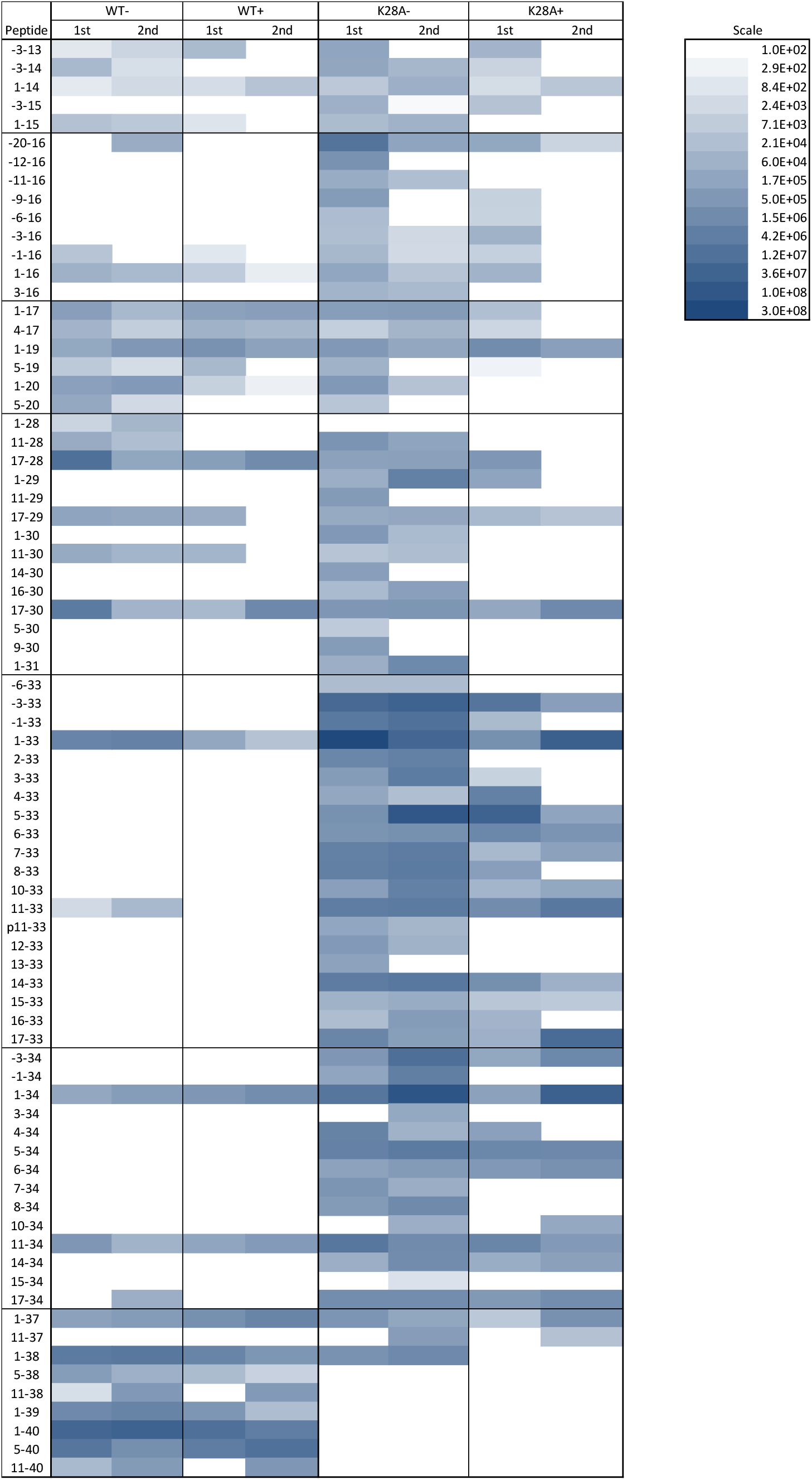
Heatmap showing APP/Aβ peptides detected using LC-ESI-MS for APP^WT^ and APP^K28A^ treated (+) and untreated (−) with cholesterol. Peptide numbering refers to the Aβ sequence, where negative numbers indicate the number of positions N-terminally of the BACE1 cleavage site. The instensity (logarithmic scale) is the peak area normalised to total protein content in the respective sample.

